# Sterols as dietary markers for *Drosophila melanogaster*

**DOI:** 10.1101/857664

**Authors:** Oskar Knittelfelder, Elodie Prince, Susanne Sales, Eric Fritzsche, Thomas Wöhner, Marko Brankatschk, Andrej Shevchenko

## Abstract

During cold acclimation fruit flies switch their feeding from yeast to plant food, however there are no robust markers to monitor it in the wild. *Drosophila melanogaster* is a sterol auxotroph and relies on dietary sterols to produce lipid membranes, lipoproteins and molting hormones. We employed shotgun lipidomics to quantify eight major food sterols in total extracts of heads, female and male genital tracts of adult flies. We found that their sterol composition is dynamic and reflective of flies diet in an organ-specific manner. Season-dependent changes observed in the organs of wild-living flies suggested that the molar ratio between yeast (ergosterol, zymosterol) and plant (sitosterol, stigmasterol) sterols is a quantifiable, generic and unequivocal marker of their feeding behavior, including cold acclimation. It provides technically simpler and more contrast readout compared to the full lipidome analysis and is suitable for ecological and environmental population-based studies.

## Introduction

*Drosophila melanogaster* (*Dm*) is an established model to study dietary and environmental impact on lipid homeostasis and development [1]. Although in the wild *Dm* is believed to feed on yeasts grown on rotten fruits and plants, it can also develop on a variety of foods, including an artificially delipidated diet supplemented with sterols [2]. *Dm* lacks Δ-6 and Δ-5 desaturases [3] and can only produce short- to medium-chain fatty acids comprising not more than one double bond [4]. However, *Dm* also makes use of dietary lipids with polyunsaturated fatty acid moieties (*e.g.* unsaturated triacylglycerols from plant oil) and incorporate them into its own lipids [5,6]. Manipulating the composition of laboratory diets helps to produce larvae and adult flies with a varying degree of unsaturation of their glycero- and glycerophospholipids and study their impact on metabolism and physiology. However, changes in lipids unsaturation span many lipid classes in a tissue-dependent manner and therefore a snapshot of the full-lipidome profile could hardly serve as a robust marker of shifting dietary preference.

Like other arthropods, *Drosophila* is a sterol auxotroph: it exclusively relies on dietary sterols to build biological membranes, lipoproteins, and to produce molting hormones [7]. Each common dietary component supplies flies with unique sterols: a commonly used yeast food (YF) is enriched in ergosterol (Erg); plant food (PF) contains phytosterols *e.g.* sitosterol (Sit), campesterol (Cam) or stigmasterol (Sti), while animal components supply cholesterol (Cho) (Figure 1). We hypothesized that, in comparison to the full lipidome composition, sterol composition could be a more specific dietary marker supporting ecological and nutritional studies also in wild-living animals.

**Figure 1.**
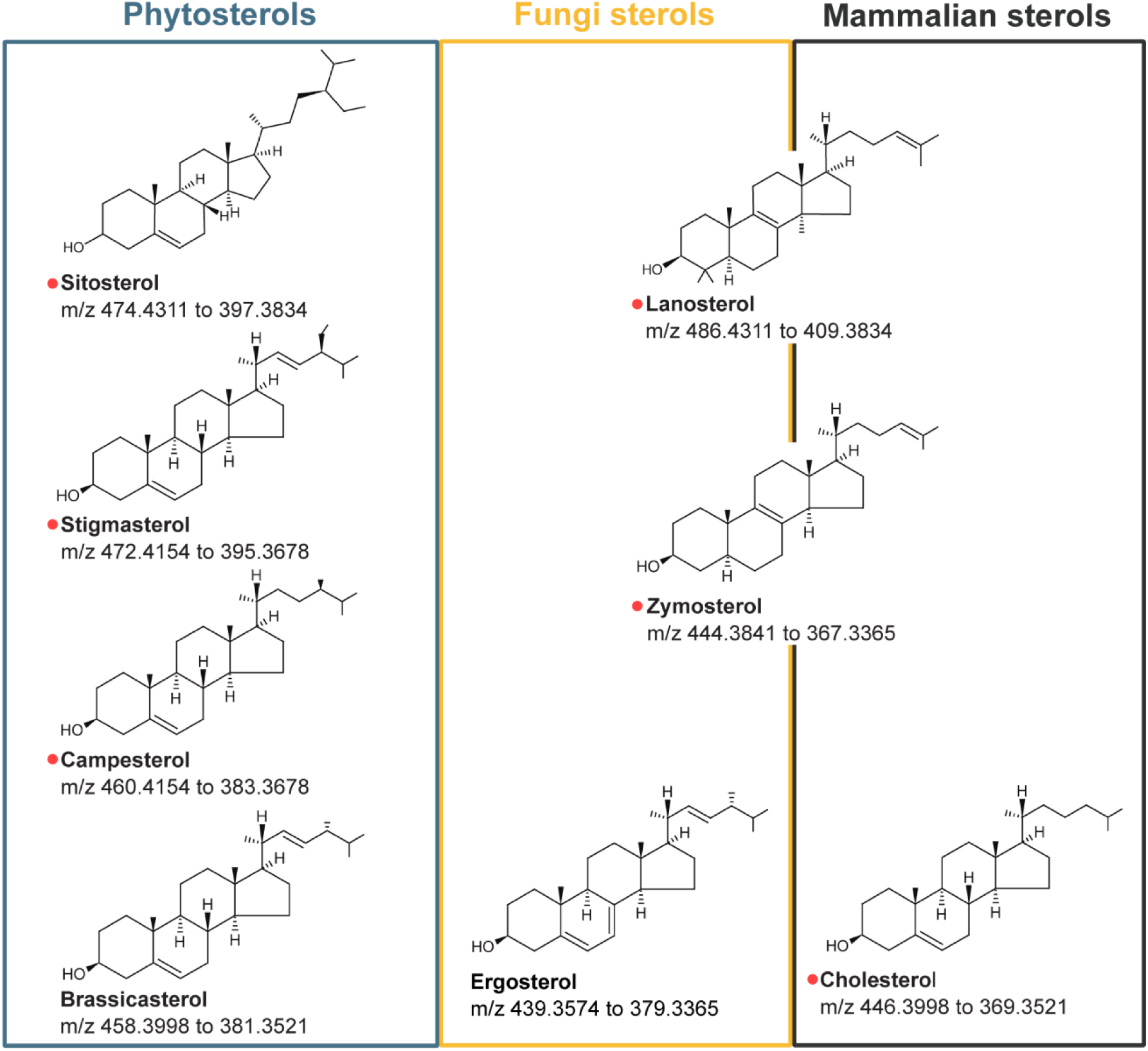
Common sterols found in fly tissues and organs [5] and grouped according to their organismal origin. Sit: sitosterol; Sti: stigmasterol; Cam: campesterol; Bra: brassicasterol; Lan: lanosterol; Zym: zymosterol; Erg: ergosterol and Cho: cholesterol. The position of Lan and Zym indicates that they are biosynthetic precursors of both Erg and Chol. SRM mass transitions used for quantifying sterols by HCD FT MS/MS is shown below their structures. Sterols with available deuterated standards are indicated by red dots.

Our recent study showed that switching from yeast to plant diet increased flies survival at low temperatures by preserving the fluidity of their biological membranes [6]. Depending on their chemical structure, sterols are differently contributing to membranes organization and fluidity [8–10]. However, these findings mostly rely on either artificial membrane vesicles or molecular dynamics calculations. It is unclear if lipidomic changes observed in the laboratory using compositionally defined foods and genetically homogeneous population of flies also play a role in the wild. It remains uncertain if the sterol composition is dynamic and reflects changes in adult flies diet that occurred within a relatively short time span. How fast are sterols exchanged and if the exchange rates is organ-specific? What sterols are preferentially incorporated by wild-living flies feeding on diverse nutrients? And, last but not least, if sterol composition follows cold acclimation in the wild?

To answer these and other important question we developed a simple and robust shotgun lipidomics method for the absolute (molar) quantification of 8 major dietary sterols in individual fly organs and monitored the dynamics of sterol exchange in adult flies reared on laboratory foods and collected in the wild.

## Results and Discussion

### Sterols quantification by shotgun lipidomics

We previously quantified the full lipidome (including 8 major sterols) of 5 fly organs by shotgun profiling [5]. To this end, total lipid extracts were split into two aliquots. One aliquot was used for quantifying glycero-, glycerophospho- and sphingolipids. Another aliquot was subjected to one-step sulfation by the sulfur trioxide - pyridine complex [5,11] followed by shotgun quantification of derivatized sterols. Sulfation increased the sensitivity and equalized the mass spectrometer response towards structurally different sterols. In this way, the full sterol complement could be quantified using a single internal standard (typically deuterated Cho) after minor adjustment of peaks abundance by individual response factors. However, despite extensive clean-up of reaction mixtures, sterols sulfation severely affected spraying stability and the method suffered from poor batch-to-batch reproducibility [12]. Also, in meantime, isotopically labeled standards of many relevant sterols have become available hence rendering the instrument response harmonization unnecessary. Therefore, instead of sulfation, we treated sterols with acetyl chloride according to Liebisch *et al*. [13], detected acetylsterols as [M+NH4]^+^ or [M+H]^+^ molecular ions, subjected them to HCD FT MS/MS and quantified using fragment ions produced by neutral loss of the acetyl group (Figure 1).

Altogether, we aimed at quantifying 8 sterols that are common to flies reared on yeast and plant laboratory foods (Figure 1) [5]. Out of those, isotopically labeled standards were available for 6 sterols, however not for the major fungi sterol Erg and the common phytosterol brassicasterol (Bra).

To quantify ergosterol, we cultivated prototrophic yeast strain W303 Y3358 on ^13^C_6_-glucose and produced fully ^13^C-labeled ergosterol [14]. In this way, we obtained a stock solution of 99% isotopically labeled ^13^C-ergosterol and then determined its molar concentration using a calibration plot made with unlabeled Erg standard (see Supplementary figure 1).

Since Bra and Sti only differ by one methyl group and also share the location of both double bonds we tested if D_6_-stigmasterol could also be used for quantifying Bra. Calibration plots of Bra and Erg covered a linear dynamic range of three orders of magnitude (0.16 to 100 µM) and were not affected by a surrogate lipid matrix (total lipid extract of bovine liver) (Supplementary Figure 1).

To quantify fly sterols we used 2/3 of a lipid material extracted from 5 heads or 5 genital tracts. The remaining extract was used for shotgun quantification of glycero- and glycerophospholipids and ceramides using the same mass spectrometry platform and software [15–18].

### Sterols in adult flies organs are exchangeable with dietary sterols

Flies incorporate sterols while growing [5], yet are they able to exchange sterols in adulthood in a fully-formed body? We monitored the sterol exchange in adult flies (5-7 days after eclosion) by swapping their diets from YF to PF and, in a separate series of experiments, from PF to YF (Figure 2).

**Figure 2.**
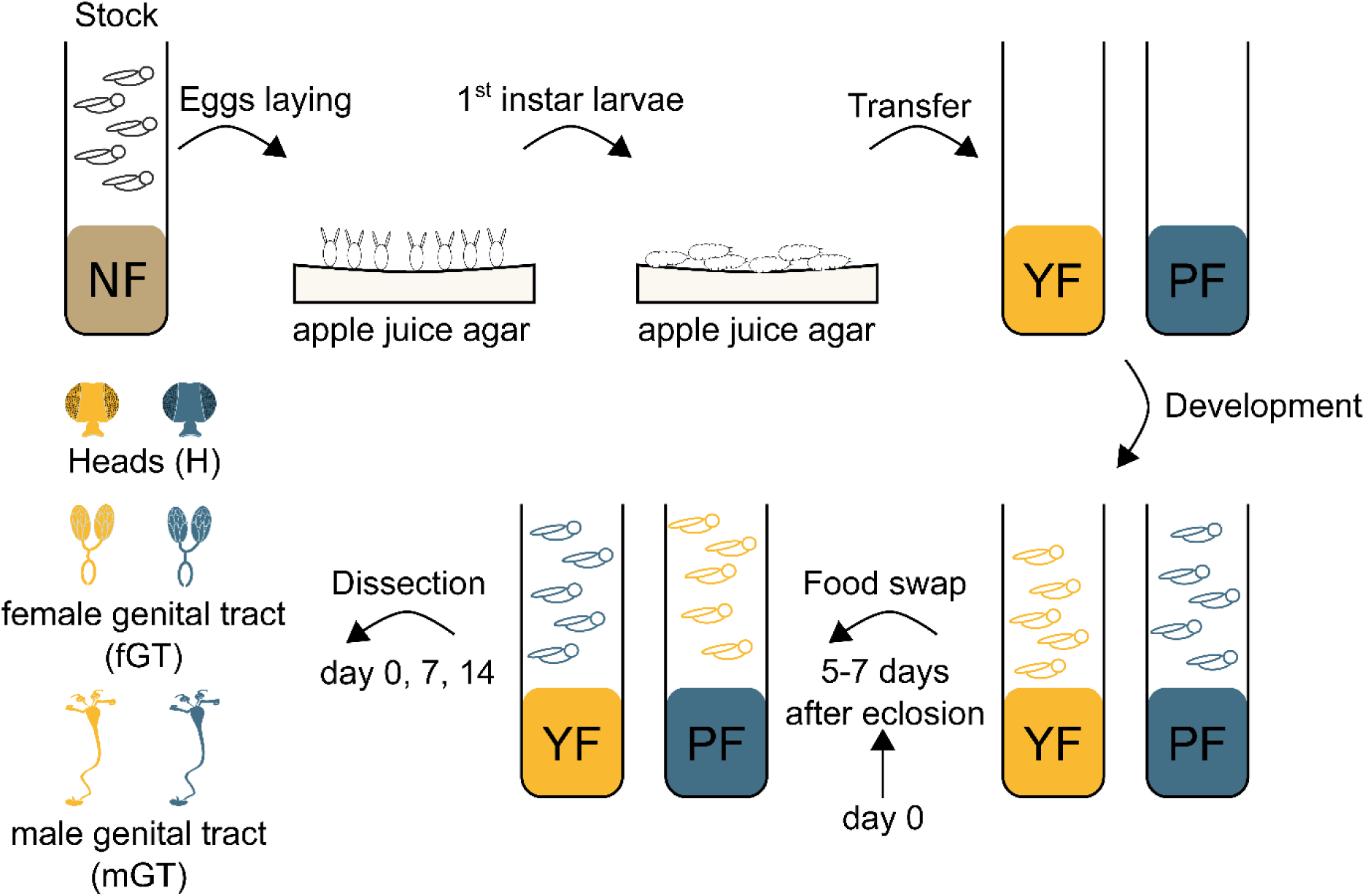
Workflow of food swapping experiments.

Flies reared on normal food (NF) containing both yeast and plant components were allowed to lay eggs on an apple juice agar plate overnight. Eggs were collected, bleached and developed to 1^st^ instar larvae on a fresh plate within one day. Larvae were collected and reared on either YF or PF till adulthood. Five to seven days after eclosion, flies were transferred to another food (PF or YF, respectively) and kept until the indicated time point (days 0, 7 and 14) (Figure 2). Heads (H), male and female genital tracts (mGT and fGT, respectively) were dissected and sterols quantified by shotgun mass spectrometry. We chose to analyze H and m/f GT to contrast the organ-dependent sterol exchange rates within the same adult animal. We reasoned that continuous sperm or eggs production in GT would require higher intake of dietary sterols compared to metabolically and morphologically conserved tissues of the fly head. At the same time, the accurate dissection of specific organs (rather than analyzing whole flies) alleviated massive compositional biases due to food lipids accumulated in the gut or fat body.

At the start (day 0) the sterol composition of H, fGT and mGT resembled the composition of nutrient sterols: Erg and Sit were major sterols in both foods and flies (Figure 3). However, relatively low abundant Erg biosynthesis precursor Zym (in flies reared on YF; for clarity further termed as YF-flies) and Cam (in PF-flies) were particularly enriched in the GTs, while at the same time PF-flies markedly rejected Sti (Figure 3). Most likely, Sit (in YF-flies) and Erg (in PF-flies) were detectable at the very low levels because of their carry-over from the NF that contained both yeast and plant material.

**Figure 3:**
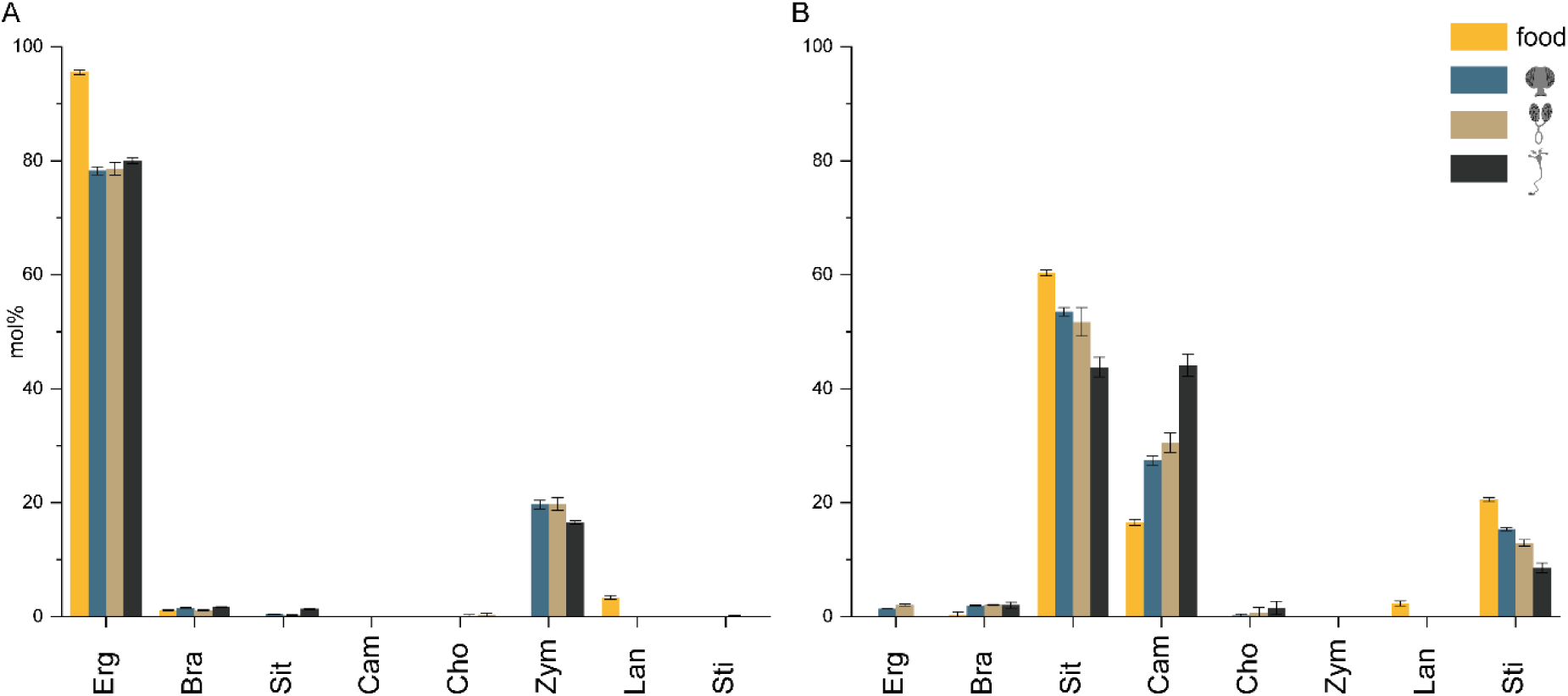
Sterol composition of yeast and plant foods and fly organs before the food swap (day 0). A. Sterol composition of yeast food (YF) and organs of flies reared on YF. B. Sterol composition of plant food (PF) and organs of flies reared on PF. Abundances of individual sterols were normalized to the total sterol content and presented in mol%; error bars indicate SD standard deviation (n = 3 with technical duplicates).

Hence, the composition of fly tissues directly followed the composition of dietary sterols, yet flies showed marked preferences towards several minor sterols. However, the relationship between the sterol abundance in food and in tissues is not direct. H and mGT of both YF- and PF-flies contained similar amount of sterols (in pmol per organ), despite YF contained 3 times more total sterols than PF (data not shown).

On the starting day prior the food swap, fGT of YF-flies contained *ca* 4-fold more sterols compared to PF-flies (Figure 4A) and 7-fold more membrane lipids (GPL and PE-Cer). This corroborated with *ca.*7-fold higher content of eggs in fGT of YF-flies that were not removed during dissection. In YF-flies both the number of eggs per ovary and the ovary area were higher than in PF-flies (Supplementary figure 2) leading to elevated fertility [6].

**Figure 4.**
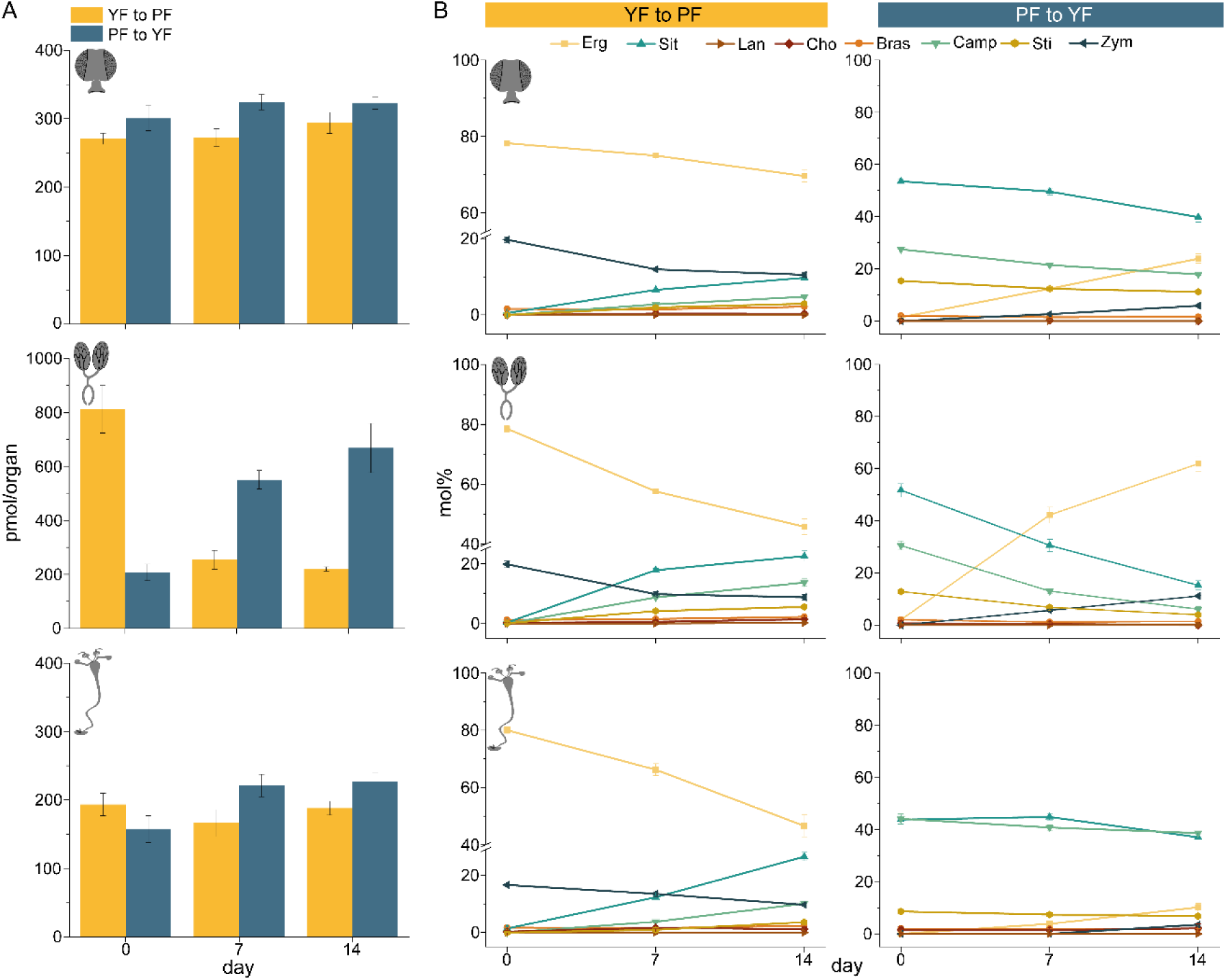
Sterol composition of fly organs during food swapping experiments (YF to PF and PF to YF) on days 0, 7 and 14. Panel A: total amount of sterols (in pmols per organ); Panel B: relative abundances (in mol%) of individual sterols (n = 3 with technical duplicates).

Within two weeks after swapping the diets (YF to PF and PF to YF), the total sterol content reached the same level as was observed in the flies reared on the substituting food. However, the sterol composition was very different. After two weeks on PF (after swapping YF to PF) only ca 35 mol% of Erg was replaced by phytosterol(s). At the same time, in the reversed direction (swapping PF to YF) Erg content increased from zero up to 60 mol% along with markedly strong depletion of both Sit and Cam.

A similar trend was observed in mGT: *ca* 30 mol% of Erg was replaced by phytosterols when changing YF to PF. However, in contrast to fGT, Erg did not replace Sit and Cam in the reversed (PF to YF) experiment.

This prompted a few interim conclusions regarding the completeness and rate of sterol exchange. Firstly, sterols are exchanged in adult flies in a tissue and sex-dependent manner at the days to weeks pace. Secondly, head sterols weakly responded to altered diets and likely reflected the composition of foods during animal growth. Thirdly, fGT sterols strongly responded to the yeast food and rapidly inreased the content of Erg. In fGT and mGT sterols profiles responded to the plant componet with approximately the same magnitude, but mGT was non-responsive to YF.

### Seasonal temperature affects the sterol composition of wild-living flies

In *Dm* cold acclimation is improved by consuming plant foods enriched with TG comprising unsaturated fatty acids [6]. Since sterol composition of tissues of adult flies is food-specific (Figure 3) and also dynamic (Figure 4), we reasoned that it could also reflect temperature-dependent dietary preferences of wild-living flies.

Wild-living adult *Dm* animals were collected in Meißen and Pirna areas (Saxony, Germany) at three different times of the year (beginning and end of August 2017 and February 2019) and their lipidomes, including sterols, were quantified by shotgun lipidomics. The sterol compositon of H, fGT and mGT was correlated with the mean temperature within the time period of two weeks prior collection (Figure 5A). Two mean temperatures (21.6°C and 18.3°C) were relatively close and we thought that corresponding fly lipidomes should reflect local variations of the wild diets, while the third (winter) collection was made at significantly lower temperature of 7.9°C.

**Figure 5:**
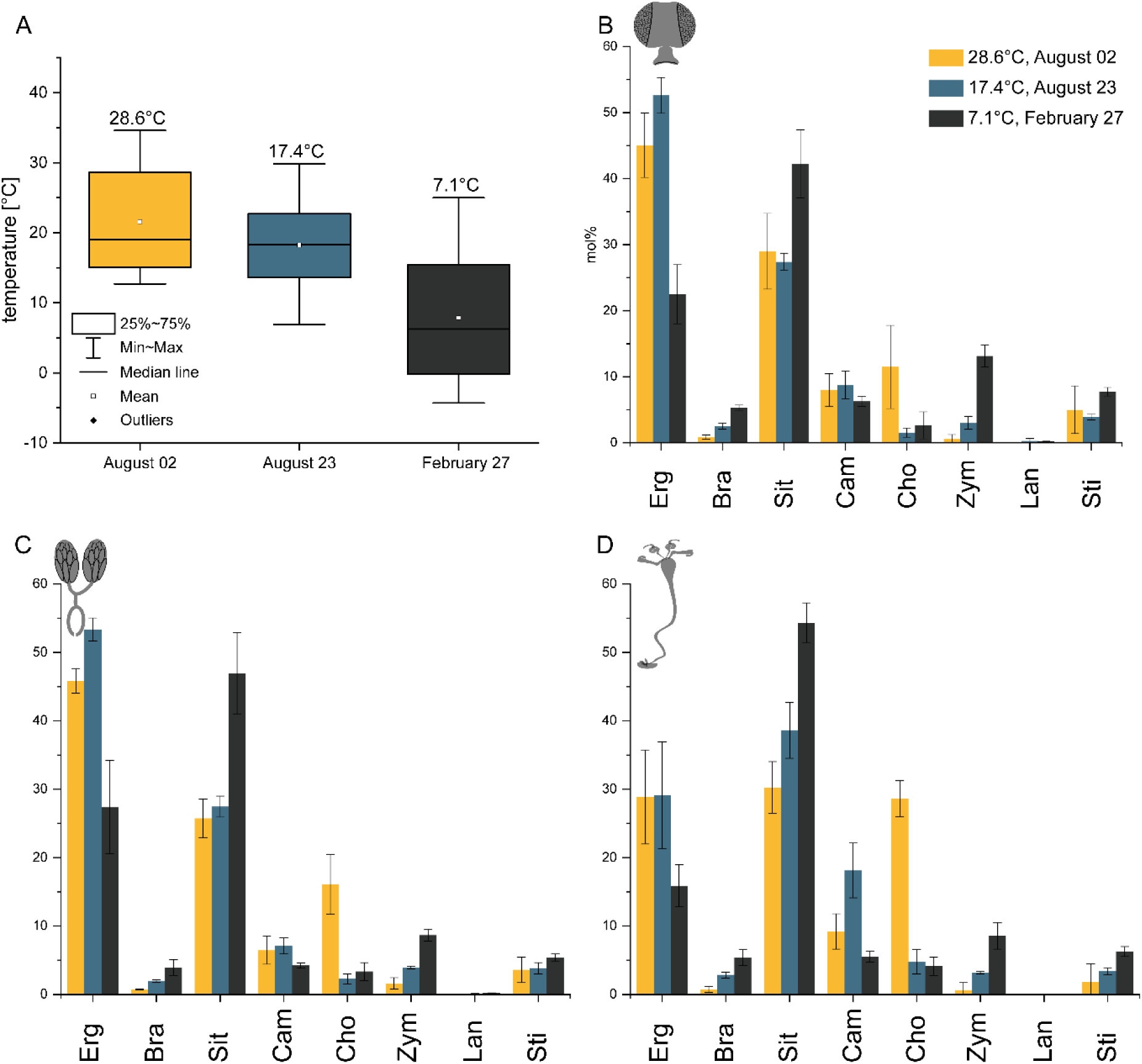
Sterol compositions of organs of wild flies collected at different temperatures. Panel A: Box plots showing temperature variation during two weeks prior the three collection times. Temperatures above the whiskers indicate collection temperatures. Panels B-D: Relative abundances (in mol%) of individual sterols in different organs at different temperatures. Collection temperatures are shown in inset. (B-D) Sterol composition of wild flies for H, fGT, and mGT, respectively (n = 3, standard deviation, technical duplicates).

**Figure 6.**
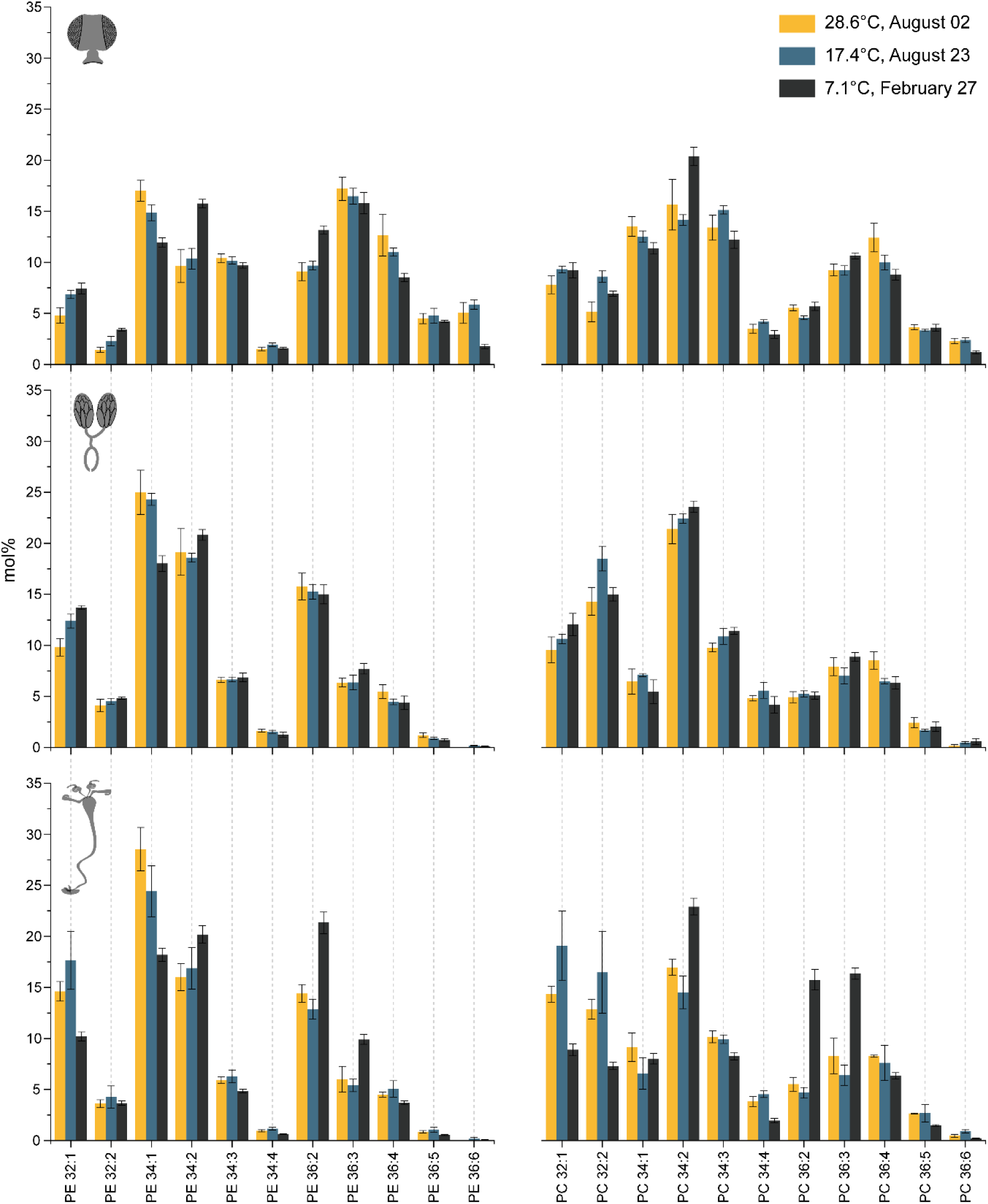
Relative abundance (in mol%) of PE and PC species in different organs of wild fles collected at different temperatures (see inset). Species annotated by lipid class and total number of carbons: total number of double bonds in the fatty acid moieties. For clarity, only species whose relative abundance exceeded 1.0 mol% are shown. (n = 3 with technical duplicates).

Sterol compositions of the three fly organs indicated that, independent of temperatures, flies were feeding on mixed diets containing comparable fractions of plant and yeast components. Interestingly, they also contained a sizable fraction of cholesterol suggesting that flies made occasional use of foods of mammalian origin [19]. Expectantly, the sterol compositon of beginning and end of August collections was similar. However, the winter flies contained approximately 50% less Erg and 50% more of Sit, compared to the summer flies, suggesting the seasonal dietary shift from yeast to plant related foods occurs in the wild (Figure 5B-D).

Interstingly, that profiles of major glycerophospholipids (PE, PC) in H and mGT of wild living animals only slightly changed with temperature. At the same time, fGT showed a clear trend toward increasing the unsaturation at lower temperature. The fly lipidome contained a substantial fraction of PC/PE species with unsaturated fatty acid moities already at the normal (21.6°C and 18.3°C) temperature. This is not suprising since other common wild living fungi *e.g. Yarrowia lipolytica* or *Pichia pastoris* could supply both ergosterol and polyunsaturated fatty acids [20–23].

Therefore, adjusting the sterol composition via dietary shift is another critical factor required for cold acclimation. It is clearly detectable also in organs whose glycerophospholipids are not strongly responding to temperature.

### Conclusions and perspectives

Among common model organisms, *Dm* is uniquely capable of building fully functional biological membranes using many dietary sterols of diverse origin and chemical structure, including mammalian sterol cholesterol, yeast ergosterol or plant sitosterol and stigmasterol, among several others. However, the biological significance and molecular basis of this structural versatility is poorly understood. We demonstrated that in mature organs of adult flies sterols were still exchangeable with food sterols. Furthermore, adjusting the sterols composition by altering the diet is an important part of the cold acclimation machinery that, at least in a few organs, functions independently of regulating membranes fluidity by changing the unsaturation of glycerophospholipids. Chol and Cam are having similar membrane ordering capacity, whereas Sit is less ordering and Sti is the least ordering from common fly sterols [8]. Hence the increase of sitosterol with concomitant depletion of (mostly) ergosterol could inrease the membrane fluidity and also contribute to cold acclimation even if unsaturated plant lipids (practically, oil-rich seeds) are not available.

In the future, it would be interesting to elucidate the consequences of sterols exchange in a broader *omics* context. In particular, an important question is if sterols could differently regulate membrane proteins/protein complexes. Together with maintaining membrane fluidity, this could be another molecular reason for remarkable tolerance of flies to the structural diversity of dietary sterols.

## Material and methods

### Fly stocks

*OregonR* stocks (#5; available from Bloomington center) were kept at 20-22°C on normal food (per liter: 22g sugar beet, 80g malt extract, 18g yeasts, 10g soy peptone, 80g cornmeal, 20g Glucose, 7g agar, 1.5g nipagin).

### Fly food recipes

Plant food (PF; calculated calories = 831kcal/L) per liter: 30g sugar beet, 45g malt extract, 2mL cold-pressed sunflower oil, 20g soy peptone, 55g cornmeal, 75g glucose, 0.7g agar, 1.5g nipagin). Yeast food (YF; calculated calories = 793 kcal/L) per liter: 20g yeast extract (Sigma Aldrich Y1625), 20g soy peptone (ROTH 2365.3), 60g glucose (ROTH X997.3), 30g sucrose, 80g yeasts, 10g agar (BioFroxx 1182GR500), 4g nipagin (ROTH 3646.2).

### Sterol swapping experiments

Embryos collected after 24h on apple-juice plates (per liter: 250mL apple-juice, 15g agar, 4g nipagin) were washed, bleached for 45sec and kept for 24h at 20-22°C on starvation plates (per liter: 250mL apple-juice, 100g glucose, 15g agar, 4g nipagin). The first-instar larvae were transferred from the starvation plates to vials containing YF or PF and kept at 19.5-20°C in a 12h/12h dark/light cycle. Once larvae reared on YF or PF reach adulthood, 5-7 days-old flies were dissected (day 0). The remaining YF- and PF-fed flies were transferred to PF and YF, respectively. Seven and 14 days after the food swap, adults were dissected (day 7 and day 14, respectively).

The adult flies were dissected in cold Phosphate-Buffered Solution (1x PBS) to remove the head and the genital tract, transferred into isopropanol on ice, snap-frozen in liquid nitrogen and stored at −80°C. For each time points and each sample, triplicates have been performed. Samples: heads (per sample: 5; male: female ratio = 2:3); male genital tracts (male gt; per sample: 5); female genital tracts (female gt; per sample: 5).

### Standards for lipid quantification

Synthetic lipid standards and Erg were purchased from Avanti Polar Lipids, Inc. (Alabaster, USA), Sigma Aldrich (Steinheim, DE) and Toronto Research Chemicals (Toronto, CA). ^13^C uniformly labeled glucose was purchased from Euriso-top (Saint Aubin, FR) and yeast nitrogen base without amino acids from Difco (LE Pont de Claix, FR). All used solvents were of at least HPLC grade. Stocks of internal standards were stored in glass ampoules at −20°C until used for the preparation of internal standard mix in 10:3 MTBE/MeOH. 700 µl internal standard mix contained: 356 pmol Cholesterol D_7_, 224 pmol Zymosterol D_5_, 215 pmol Campesterol D_6_, 207 pmol Sitosterol D_6_, 201 pmol Lanosterol D_6_, 418 pmol Stigmasterol D_6_, 233 pmol Desmosterol D_6_, 196 pmol ^13^C Ergosterol, 417 pmol 50:0 TG D_5_, 116 pmol 34:0 DG D_5_, 220 pmol 25:0 PC, 77 pmol LPC, 107 pmol 25:0 PS, 354 pmol 25:0 PE, 85 pmol 13:0 LPE, 96 pmol 25:0 PI, 109 pmol 25:0 PG, 145 pmol 30:1 Cer, 91 pmol 25:0 PA, 45 pmol 13:0 LPA, 178 pmol 29:1 CerPE, 38 pmol 13:0 LPI, 54 pmol 56:4 CL, 59 pmol 13:0 LPS, 75 pmol 13:0 LPG.

^13^C uniformly labeled Erg was produced in the prototrophic yeast strain W303 Y3358. A single colony was inoculated in 25 ml sterile filtered synthetic defined medium (20 g/l glucose, 6.7 g/l yeast nitrogen base without amino acids). Yeast was incubated for 21 h at 30°C to reach stationary phase and then pelleted by centrifugation (10 min, 4000 g). The medium was removed and the pellet was washed twice with 1 ml H_2_O. The corresponding pellet was split into 6 samples. These pellets were reconstituted in 300 µl isopropanol (IPA) each and homogenized with 0.5 mm zirconia beads on a tissuelyser II for 20 min at 30 Hz. The dried homogenates were saponified with 1 ml 3% KOH in methanol for 2 h at 90°C each. Ergosterol was extracted twice with 2 ml hexane and 1 ml H_2_O and the combined organic phases from all samples were evaporated. Finally, Erg was reconstituted in 1 ml MeOH and stored at −20 C. ^13^C-ergosterol was quantified by adding a known amount of ^12^C-ergosterol and performing parallel reaction monitoring.

### Lipid extraction and quantification by shotgun mass spectrometry

5 Dm heads (fixed ratio of 2 male heads and 3 female heads) or 5 genital tracts of corresponding sex were homogenized with 1 mm zirconia beads in a cooled tissuelyzer for 2 x 5 min at 30 Hz in 200 µl IPA. The homogenate was evaporated in a vacuum desiccator to complete dryness. For YF and PF, 200 – 300 mg wet weight were homogenized in 600 µl IPA with 1 mm zirconia beads in a cooled tissuelyzer for 20 min at 30 Hz. An aliquot corresponding to 4 mg food wet weight was used for lipid extraction. All samples were extracted according to [24]. In brief, 700 µl internal standard mix in 10:3 MTBE/MeOH was added to each sample and vortexed for 1 h at 4°C. After the addition of 140 µl H_2_O, samples were vortexed for another 15 min. Phase separation was induced by centrifugation at 13200 rpm for 15 min. The organic phase was transferred to a glass vial and evaporated. Samples were reconstituted in 300 µl 1:2 MeOH/CHCl_3_. 100 µl were transferred to a new vial and used for lipidome analysis. To quantify sterols 200 µl of lipid extract were evaporated and acetylated with 300 µl 2:1 CHCl_3_/acetyl chloride for 1 h at room temperature (modified from Liebisch et al., 2006). After evaporation sterol samples were reconstituted in 200 µl 4:2:1 IPA/MeOH/CHCl_3_ + 7.5 mM ammonium formate (spray solution). For sterol measurements, samples were 1:5 diluted with spray solution. For lipidome measurements, samples were 1:10 diluted with spray solution.

Mass spectrometric analysis was performed on a Q Exactive instrument (Thermo Fischer Scientific, Bremen, DE) equipped with a robotic nanoflow ion source TriVersa NanoMate (Advion BioSciences, Ithaca, USA) using nanoelectrospray chips with a diameter of 4.1 µm. The ion source was controlled by the Chipsoft 8.3.1 software (Advion BioSciences). Ionization voltage was + 0.96 kV in positive and − 0.96 kV in negative mode; backpressure was set at 1.25 psi in both modes. Samples were analyzed by polarity switching [25]. The temperature of the ion transfer capillary was 200 °C; S-lens RF level was set to 50%. All samples were analyzed for 10 min. FT MS spectra were acquired within the range of m/z 400– 1000 from 0 min to 0.2 min in positive and within the range of m/z 350–1000 from 5.2 min to 5.4 min in negative mode at the mass resolution of R m/z 200 = 140000; automated gain control (AGC) of 3×10^6^ and with the maximal injection time of 3000 ms. *t*-SIM in positive (0.2 to 5 min) and negative (5.4 to 10 min) mode was acquired with R m/z 200 = 140000; automated gain control of 5 × 10^4^; maximum injection time of 650 ms; isolation window of 20 Th and scan range of m/z 400 to 1000 in positive and m/z 350 to 1000 in negative mode, respectively. The inclusion list of masses targeted in *t*-SIM analyses started at m/z 355 in negative and m/z 405 in positive ion mode and other masses were computed by adding 10 Th increment (i.e. m/z 355, 365, 375, …) up to m/z 1005. Acetylated sterols were quantified by parallel reaction monitoring (PRM) FT MS/MS in an additional measurement. FT MS spectra within the range of m/z 350-1000 were acquired from 0 min to 0.2 min and *t*-SIM ranging from m/z 350 to 500 were acquired from 0.2 min to 4 min with the same settings as described above. PRM spectra were acquired from 4 min to 10 min. For PRM micro scans were set to 1, isolation window to 0.8 Da, normalized collision energy to 12.5%, AGC to 5 × 10^4^ and maximum injection time to 3000 ms. All spectra were pre-processed using repetition rate filtering software PeakStrainer [26] and stitched together by an in-house developed script [17]. Lipids were identified by LipidXplorer software [27]. Molecular Fragmentation Query Language (MFQL) queries were compiled for acetylated sterols, PA, LPA PC, PC O-, LPC, LPC O-, PE, PE O-, LPE, PI, LPI, PS, LPS, PG, LPG, CL, CerPE, Cer, TG, DG lipid classes. The identification relied on accurately determined intact lipid masses (mass accuracy better than 5 ppm). Lipids were quantified by comparing the isotopically corrected abundances of their molecular ions with the abundances of internal standards of the same lipid class. For acetylated sterols, the specific fragment (loss of acetyl group) was used for quantification.

## Supporting information

Sterols as dietary markers for Drosophila melanogaster supplement

## Acknowledgments

We are greatly indebt to our late colleague Dr.Suzanne Eaton who should have been a rightful co-authors of this manuscript. Dr.Eaton inspired us to work on the lipidomics of temperature acclimation and enormously contributed to designing and organizing the study. Due to a tragic loss of life she has never got a chance to see the results she envisioned.

We also acknowledge the Simon Alberti laboratory for supplying the prototrophic yeast strain W303 Y3358 and Aliona Bogdanova from the Protein Expression Facility at MPI Dresden for her help with yeast cultivation. We thank the members of Shevchenko lab for their help and support and Clara Poupault for useful comments and proofreading. Work in AS and MB laboratory was supported by FOR 2682 grant from Deutsche Forschungsgemeinschaft (DFG).

