## Supplementary material for "Sterols as dietary markers for *Drosophila melanogaster*": Sterols as dietary markers for Drosophila melanogaster supplement

Supplementary figures

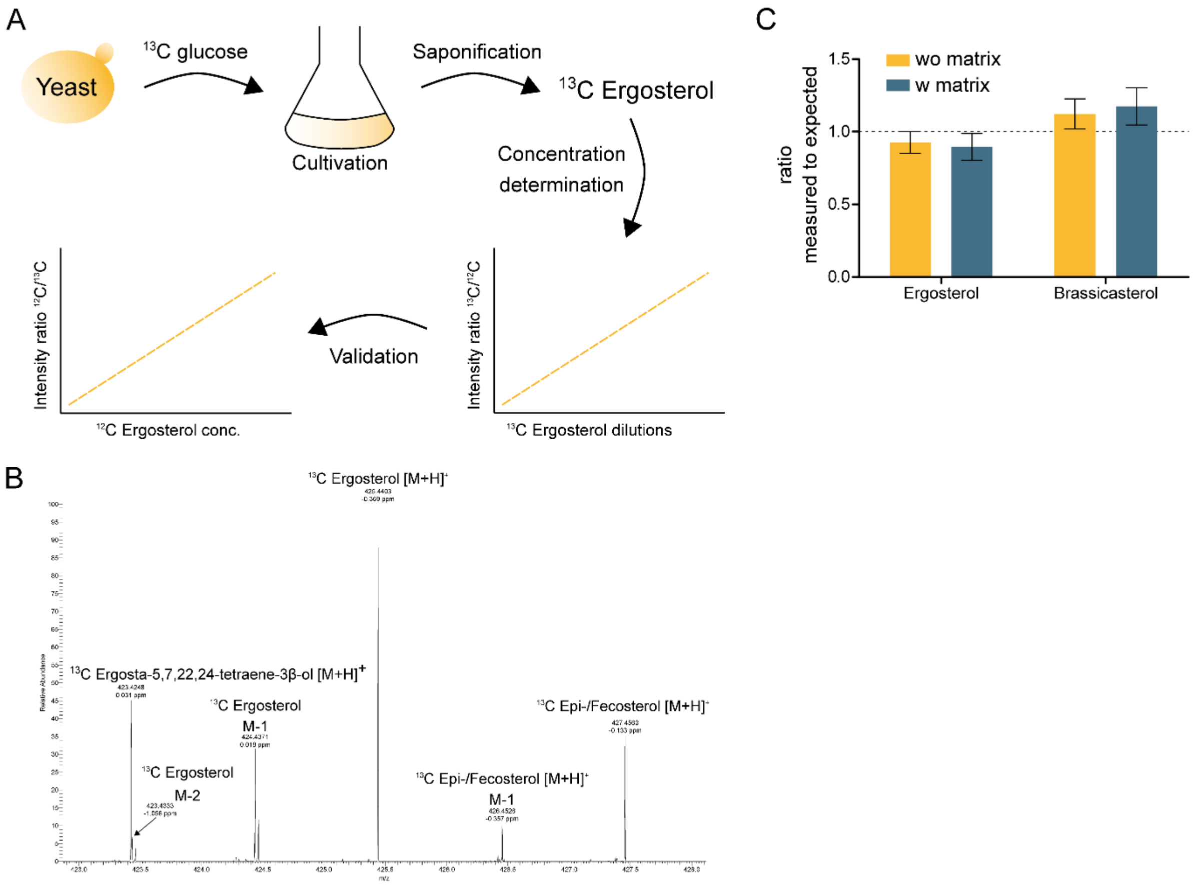


Supplementary figure 1: Production and validation of $^{13}$C ergosterol and quantification of brassicasterol by stigmasterol $d_6$. A. Workflow of $^{13}$C ergosterol production in yeast. The prototrophic yeast strain W303 Y3358 was cultivated with uniformly labeled $^{13}$C glucose as the sole carbon source. After reaching stationary phase, the yeast pellet was homogenized and subjected to saponification. $^{13}$C ergosterol was extracted and quantified by $^{12}$C ergosterol. To validate the result, $^{13}$C ergosterol was used to quantify known amounts of $^{12}$C ergosterol. B. Mass spectrum of non-derivatized $^{13}$C ergosterol including labeled ergosterol precursors. C. Validation of ergosterol and brassicasterol quantification. Calibration curves with concentrations ranging from 0.16 µM to 100 µM in absence or presence of lipid matrix (total lipid extract of bovine liver) were determined with the corresponding internal standards ($^{13}$C ergosterol and stigmasterol $d_6$) and compared to the expected concentrations. Results are presented as measured to expected ratios.

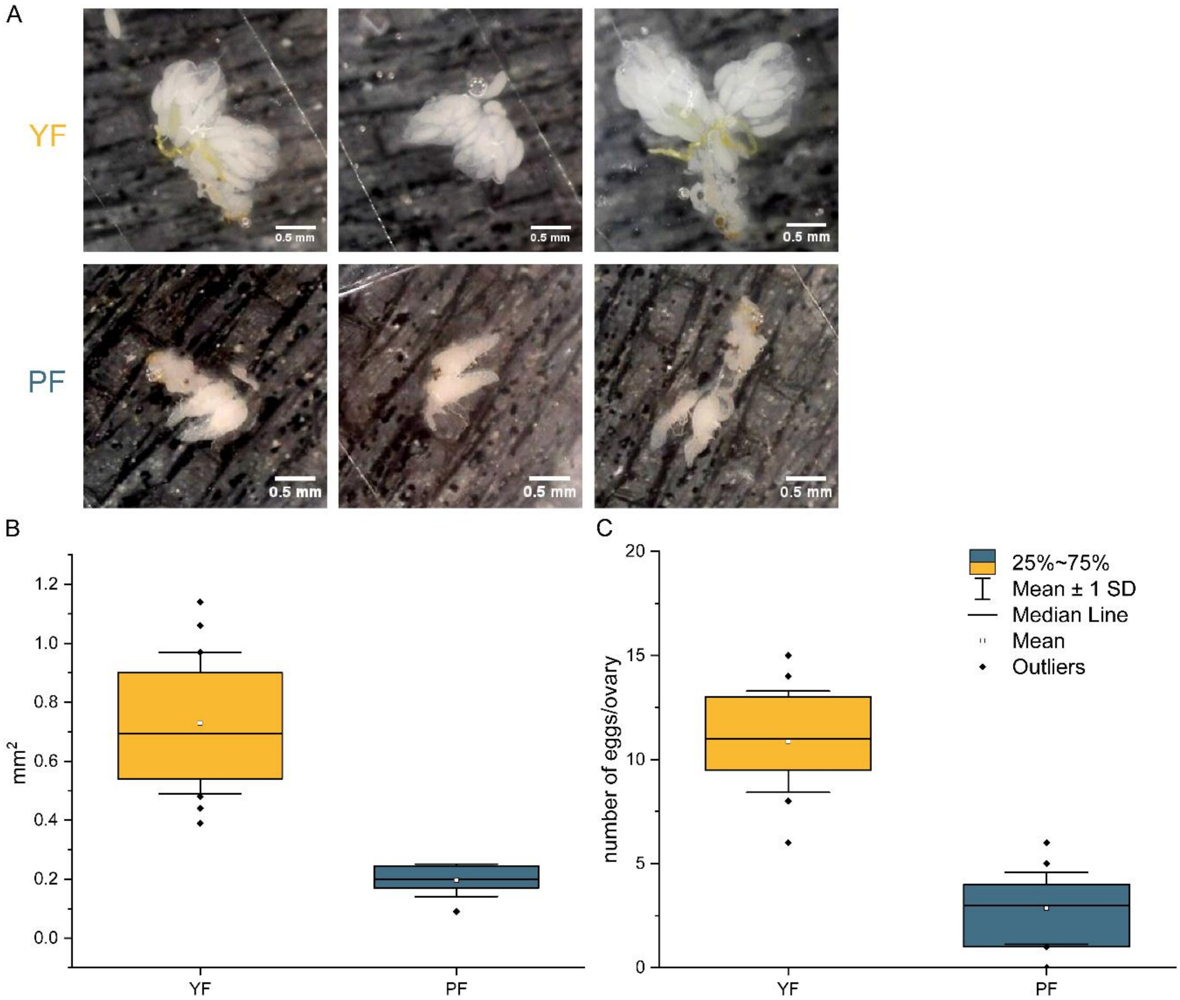


Supplementary figure 2: Influence of diet on female genital tract size and number of eggs per ovary. A. Representative images of dissected female genital tracts from flies either raised on yeast (YF) or plant food (PF) are depicted. B. Comparison of the ovary area in $mm^2$ between the two different diets (YF: 6 flies, 2 ovaries, PF: 4 flies, 2 ovaries). C. Obtained ovaries were further dissected and the number of eggs per ovary was determined (YF: 10 flies, 2 ovaries, PF: 10 flies, 2 ovaries).
